# Efficient and Modular CRISPR-Cas9 Vector System for *Physcomitrella patens*

**DOI:** 10.1101/674481

**Authors:** Darren R. Mallett, Mingqin Chang, Xiaohang Cheng, Magdalena Bezanilla

**Affiliations:** Department of Biological Sciences, Dartmouth College, Hanover NH 03755; Plant Biology Graduate Program, University of Massachusetts, Amherst 01002

**Author notes:** Authors contributed equally (Darren R. Mallett, Mingqin Chang). Corresponding author., M.B. Emails: D.R.M., M.C., X.C.

**Keywords:** P. patens, CRISPR, vector system, genome editing, homology-directed repair, multiplexing

## Abstract

CRISPR-Cas9 has been shown to be a valuable tool in recent years, allowing researchers to precisely edit the genome using an RNA-guided nuclease to initiate double-strand breaks. Until recently, classical RAD51-mediated homologous recombination has been a powerful tool for gene targeting in the moss *Physcomitrella patens*. However, CRISPR-Cas9 mediated genome editing in *P. patens* was shown to be more efficient than traditional homologous recombination (Collonnier et al. 2017). CRISPR-Cas9 provides the opportunity to efficiently edit the genome at multiple loci as well as integrate sequences at precise locations in the genome using a simple transient transformation. To fully take advantage of CRISPR-Cas9 genome editing in *P. patens*, here we describe the generation and use of a flexible and modular CRISPR-Cas9 vector system. Without the need for gene synthesis, this vector system enables editing of up to 12 loci simultaneously. Using this system, we generated multiple lines that had null alleles at four distant loci. We also found that targeting multiple sites within a single locus can produce larger deletions, but the success of this depends on individual protospacers. To take advantage of homology-directed repair, we developed modular vectors to rapidly generate DNA donor plasmids to efficiently introduce DNA sequences encoding for fluorescent proteins at the 5’ and 3’ ends of gene coding regions. With regards to homology-directed repair experiments, we found that if the protospacer sequence remains on the DNA donor plasmid, then Cas9 cleaves the plasmid target as well as the genomic target. This can reduce the efficiency of introducing sequences into the genome. Furthermore, to ensure the generation of a null allele near the Cas9 cleavage site, we generated a homology plasmid harboring a “stop codon cassette” with down-stream near-effortless genotyping.

## INTRODUCTION

Recent advances in genome editing are actively revolutionizing the fields of genetics, biotechnology, medicine, and agronomics. The creation of double-strand breaks in DNA by site-specific nucleases is an essential step in efficient genome editing. In the last few years, CRISPR has been employed in a wide array of organisms to create double-strand breaks with great success, and the design is quite straightforward (Sander and Joung 2014). Adopted from type II CRISPR systems in bacteria, introduction of the Cas9 enzyme and a programmable single-guide RNA (sgRNA) into cells is among the most common to generate controlled DNA double-strand breaks (Makarova et al. 2011; Jinek et al. 2012; Sander and Joung 2014; Malzahn, Lowder, and Qi 2017). The sgRNA contains ∼20 bases homologous to a DNA target of interest at the 5’ end (known as the protospacer) and a region that binds the Cas9 nuclease (Jinek et al. 2012; Nishimasu et al. 2014). The Cas9:sgRNA complex binds and cleaves a target DNA sequence if the protospacer is directly 5’ to a protospacer adjacent motif (PAM) on the noncomplementary DNA strand (Jinek et al. 2012; Nishimasu et al. 2014).

Double-strand breaks, which are a form of DNA damage, are repaired by one of two major pathways: non-homologous end joining or homology-directed repair (also referred to as homologous recombination) (Sander and Joung 2014; Moynahan and Jasin 2010; Chang et al. 2017). Double-strand breaks repaired by non-homologous end joining often result in inserted or deleted nucleotides (“indels”), especially when microhomology-mediated end joining (also known as alternative end joining) is employed (Chang et al. 2017). The resulting indels often disrupt gene function by potentially altering the translational reading frame within a protein coding region. Alternatively, homology-directed repair uses a DNA template that shares homology with both sides of the break to accurately repair the DNA (Moynahan and Jasin 2010). By taking advantage of this pathway, it is possible to precisely alter a gene of interest by providing a DNA “donor” template containing desired modifications together with Cas9 and the sgRNA. However, the overall activity of homology-directed repair is quite low in a variety of organisms, and thus the majority of double-strand breaks are repaired by non-homologous end joining (Sargent, Brenneman, and Wilson 1997; Beucher et al. 2009; Puchta 2005).

In plants, CRISPR has been used to perform a variety of gene modifications, including the targeted mutagenesis of genes related to crop yield (M. Li et al. 2016) as well as host genes required for disease pathogenesis (Wang et al. 2016; Pyott, Sheehan, and Molnar 2016). Successful editing events have been reported in rice (Shan et al. 2013), maize (Liang et al. 2014), and *Arabidopsis thaliana* (J.-F. Li et al. 2013; Feng et al. 2014), as well as in polyploid crops deemed difficult for gene editing such as strawberry (F. M. Wilson et al. 2019), wheat (Zhang et al. 2016), and cotton (C. Li, Unver, and Zhang 2017; Chen et al. 2017), among many others (see (Malzahn, Lowder, and Qi 2017) for a review). The majority of these gene editing experiments resulted in gene knock-out via non-homologous end joining. Gene targeting using homology-directed repair has been quite challenging in seed plants, with most attempts reporting low success rates (J.-F. Li et al. 2013; Svitashev et al. 2015; Shi et al. 2017; Svitashev et al. 2016; Butler et al. 2016; Čermák et al. 2015; Schiml, Fauser, and Puchta 2014). The model moss *Physcomitrella patens* has been used over the last few decades to study various fundamental processes of plant biology. *P. patens* is known to be exceptionally amenable to genetic manipulation due to its ability to perform high rates of homologous recombination, especially when linear DNA is supplied (Schaefer and Zryd 1997; Prigge and Bezanilla 2010; Kamisugi and Cuming 2009). Recently, both CRISPR-mediated gene knock-out (using non-homologous end joining) (Lopez-Obando et al. 2016) and gene knock-in (using homology-directed repair) (Collonnier et al. 2017) have high rates of success in *P. patens*, including the ability to target multiple genes in a single, transient transformation (i.e. “multiplexing”) (Lopez-Obando et al. 2016).

Here we describe an efficient and modular CRISPR vector system for use in *Physcomitrella patens*. In this system, protospacer sequences are synthesized as oligonucleotides and are efficiently ligated into entry vectors containing the sgRNA expression cassette, eliminating the need for gene synthesis. Using Multisite Gateway cloning (Invitrogen), multiple entry vectors recombine with a single destination vector containing Cas9 for efficient multiplexing. By co-transforming three expression vectors with different antibiotic selection cassettes, it is possible to target up to 12 genomic sites in a single, transient transformation. Here we showcase the multiplexing capabilities of the system by simultaneously targeting six genes, as well as creating gene deletions using adjacent sgRNAs. In addition, we describe a simple, yet effective way to clone homology fragments flanking genes encoding fluorescent proteins to efficiently generate donor template DNA for use with homology-directed repair. Likewise, we introduce a novel concept to use homology-directed repair to knock-in DNA encoding for multiple stop codons in each reading frame to allow for a controlled gene knock-out with near-effortless genotyping. Lastly, we explore the issues that arise when donor template DNA contains the protospacer sequence during homology-directed repair experiments and describe ways to avoid these issues.

## RESULTS

### Optimal expression of the sgRNA results in high efficiency editing

To test if RNA polymerase III promoters from other organisms could be used in moss, we compared the efficiency of genome editing using either a rice U3 or a *P. patens* U6 promoter to drive expression of the guide RNA. As a rapid visual test for genome editing, we designed a protospacer that targets the 5’ end of the coding sequence of green fluorescent protein (GFP) (Figure 1A). The final Cas9 expression plasmids harboring either the PpU6 or the OsU3 promoter were then transformed separately into NLS-4, a moss line that stably expresses nuclear-localized GFP fused to β-glucuronidase (GUS) (Bezanilla, Pan, and Quatrano 2003). Thus, plants lacking nuclear green fluorescence after transformation represent genome editing events leading to a loss-of-function mutation in the GFP:GUS fusion protein.

**Figure 1.**
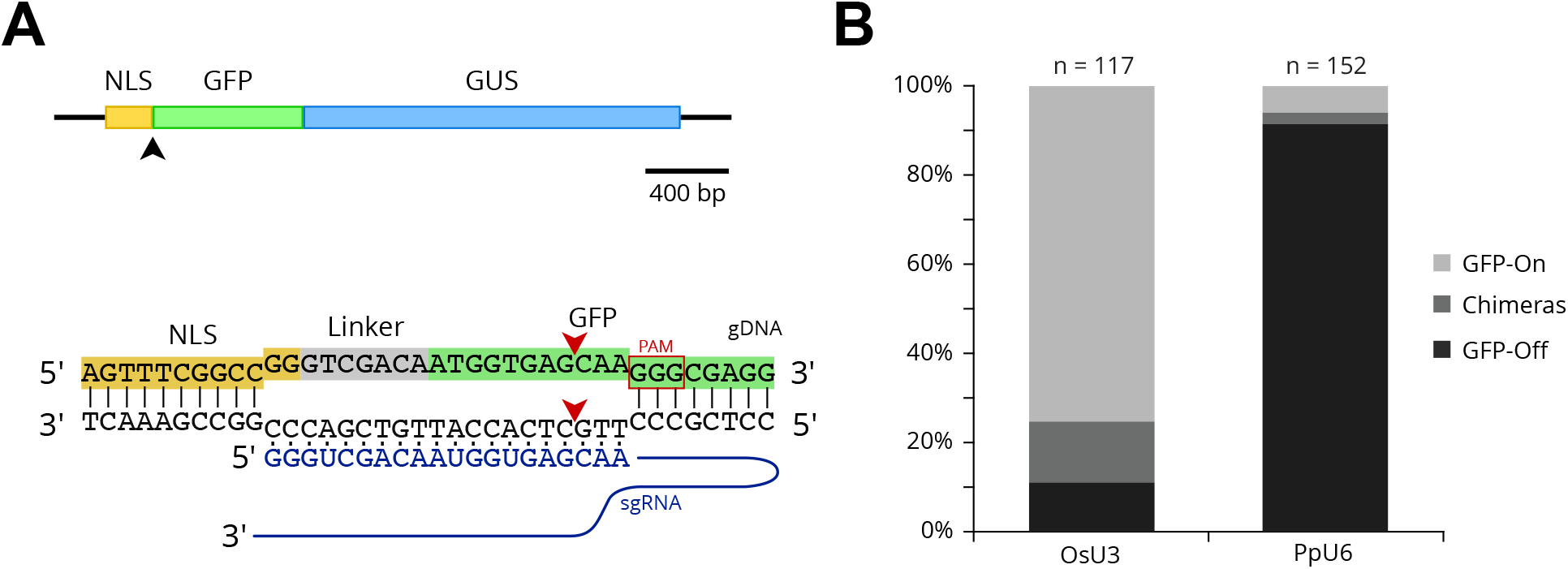
(A) A gene model of the NLS-GFP-GUS reporter. The black arrowhead indicates the expected Cas9 cleavage site. Below the gene model, the DNA sequence at the junction of the NLS and GFP gene fragments is shown with the sgRNA binding site. Red arrowheads indicate the expected cleavage site by Cas9 at the 5’ end of the GFP coding region. The PAM sequence is indicated with a red box. (B) A stacked bar graph representing the efficiency of editing the NLS-GFP-GUS gene using either OsU3 or the PpU6 promoters to drive expression of the sgRNA. Plants expressing the reporter (GFP-on) are most likely not edited. Plants lacking NLS-GFP-GUS expression (GFP-off) are edited. Chimeras indicate plants with GFP-GUS expression in only a portion of the plant.

After two weeks on selection, we imaged transformed plants using fluorescence microscopy to visualize the presence (or absence) of nuclear fluorescence. We found that plants transformed with the OsU3::sgRNA resulted in 11.1% of plants lacking GFP signal. In comparison, 91.4% of plants transformed with the PpU6::sgRNA lacked GFP signal (Figure 1B). We verified that NLS-GFP-GUS was edited in a subset of plants lacking GFP signal via Sanger sequencing (*n* = 10, OsU3::sgRNA transformants, Supplemental Figure 1). Additionally, we observed that 13.7% of plants transformed with OsU3::sgRNA contained nuclear fluorescence in a portion of the plant, giving rise to a population of chimeric plants. Interestingly, of the plants transformed with PpU6::sgRNA, we only observed chimeras in 2.6% of the plants (Figure 1B). We reasoned that chimeras arise as a result of the Cas9 nuclease cleaving the NLS-GFP-GUS reporter after the initial cell division of the transformed protoplast. Given that the OsU3 promoter resulted in fewer plants lacking GFP and a larger percentage of chimeric plants as compared to the PpU6 promoter, our results suggest that the PpU6 promoter is more efficient at expressing the sgRNA in moss protoplasts.

### Flexible vector system enables simultaneous targeting of multiple genomic sites

Here we present the development of a vector system that rapidly and flexibly allows for simultaneous targeting of multiple genomic sites. Our system builds on vectors developed for rice (Miao et al. 2013) with modifications and enhancements to increase expression and transformation efficiency in *P. patens*. In the moss vector system, the sgRNA expression cassette resides in a Gateway entry vector (Invitrogen) and consists of the PpU6 promoter followed by DNA encoding the sgRNA (Figure 2A). The protospacer sequence is easily modified to target a specific genomic locus. Two reverse-complementary oligonucleotides containing a custom protospacer sequence are annealed together and directionally ligated into an entry vector that has been linearized by two *BsaI* sites (Figure 2B). Using site-specific recombination, the ligated entry vector recombines with a destination vector containing the Cas9 expression cassette in which Cas9 expression is driven by the maize ubiquitin promoter to create a final expression vector containing both components. We chose to use the maize ubiquitin promoter as this is a well-documented promoter for high-level expression in moss (Bezanilla, Pan, and Quatrano 2003; Saidi et al. 2005).

**Figure 2.**
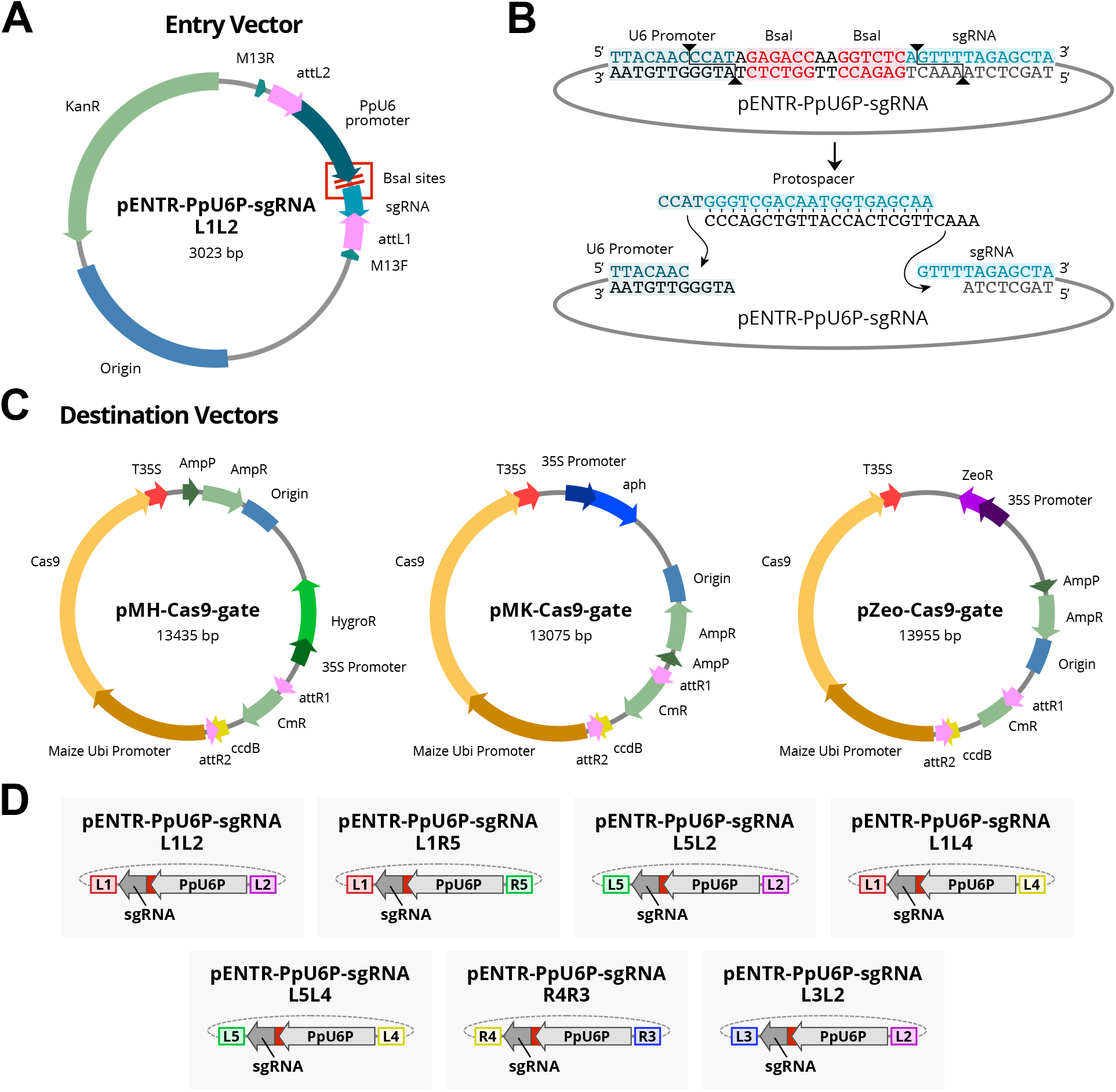
(A) A plasmid map of the entry vector containing the U6 promoter to drive expression of the sgRNA flanked by Gateway att sites. (B) The DNA sequence of U6 promoter-sgRNA junction within the entry clone separated by inverted BsaI sites. Upon digestion of the entry clone with BsaI, unique vector overhangs allow for directional ligation of custom oligonucleotides containing the protospacer sequence. (C) Plasmid maps of the destination vectors pMH-Cas9-gate, pMK-Cas9-gate, and pZeo-Cas9-gate for hygromycin, G418, and Zeocin selection in plants, respectively. (D) Entry vectors are shown that were generated based on the plasmid shown in (A) with modified att sites enabling compatibility with Multisite Gateway reactions.

For maximum flexibility, we generated three destination vectors comprising different antibiotic resistance genes for selection in plants (Figure 2C). To target multiple genomic sites in one transformation, we took advantage of Multisite Gateway (Invitrogen), which enables directional stitching of up to four DNA fragments. We generated Multisite Gateway entry vectors enabling construction of a single expression vector that expresses Cas9 and up to four unique sgRNAs simultaneously (Figure 2D). Using this powerful system, it is possible to target up to 12 different genomic sites using each of the three destination vectors upon simultaneous selection with hygromycin, G418, and Zeocin.

### Targeting multiple, distant genomic sites

To test successful targeting of multiple genomic sites in one transformation, we designed protospacer oligos to target six genomic sites (site 1: Pp3c8_18830V3.1; site 2: Pp3c18_4770V3; site 3: Pp3c22_15110V3; site 4: Pp3c4_16430V3; site 5: Pp3c8_18850V3; site 6: Pp3c23_15670V3). Of these sites, two of them (site #1 and site #5) have targeting sites 8727 bp apart on the same chromosome and the remaining four have targeting sites on different chromosomes. We generated two expression vectors, each harboring three sgRNA expression cassettes (Figure 3A). Protoplasts were subsequently co-transformed and selected for both expression constructs. We subsequently genotyped the remaining plants using T7 endonuclease, an enzyme that recognizes and cleaves regions of mismatching bases present in dsDNA. Thus, indel mutations can be easily detected when the DNA to be tested is annealed to wild type DNA. To start, we screened 24 plants at site #1 and site #4. In these 24 plants, we were unable to obtain T7 cleavage at site #1. However, 15 plants resulted in T7 cleavage at site #4. We sequenced these plants and confirmed the presence of indel mutations in all 15 plants, 6 plants of which contained frameshift mutations (Figure 3B). We subsequently screened the remaining sites (sites #2, #3, #5, and #6) in these 6 plants and found edits at sites #3, #5, and #6, but not at site #2 (Figure 3B; Supplementary Figure 2). Taken together, these results provide evidence that sgRNA expression from two separate vectors enables targeting of multiple genes in a single transformation event.

**Figure 3.**
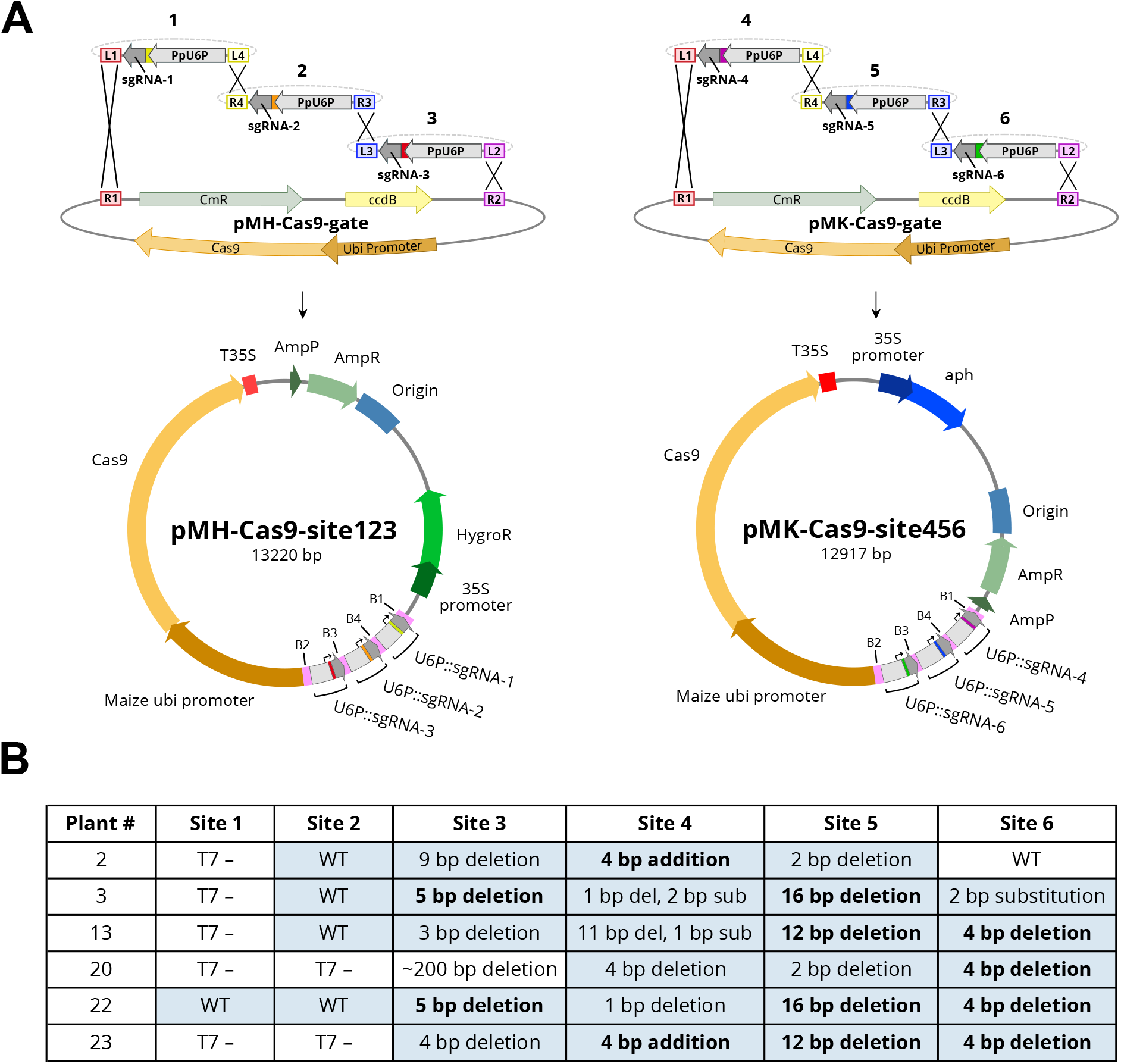
(A) A schematic showing the directional Multisite Gateway LR recombination reactions between entry clones containing the sgRNA expression cassette and each destination vector to create two final expression constructs (final plasmid maps are drawn to scale). (B) A table showing the genotyping results of 6 plants regenerated from protoplasts that were transformed with both plasmids simultaneously. Plants were initially screened by a T7 endonuclease assay. “T7-” indicates that T7 endonuclease was unable to recognize a mismatch when transformant gDNA was paired with WT gDNA. Blue-shaded cells indicate sequencing was performed on that site. Bolded sequencing results with identical values in the same column indicate identical edits at that site. For sequencing data, see Supplemental Figure 2.

### Targeting adjacent genomic sites can result in large deletions

The ability to target a region with multiple sgRNAs has been shown to be beneficial in creating knock-out mutations and large deletions in other model systems, including soybean (Cai et al. 2018), human cells (He et al. 2015), zebrafish (Xiao et al. 2013), and yeast (Hao et al. 2016). Protospacer sequences have variable probabilities of producing out-of-frame mutations, depending on the surrounding microhomology in the genomic DNA available for microhomology-mediated end joining repair (Bae et al. 2014). Additionally, an indel that would cause a frameshift mutation in a protein-coding gene may not disrupt the function of non-coding DNA. To increase the chances of making knock-out mutations and easily visualizing them on a gel, it may be beneficial to excise a region of the gene using two sgRNAs. To test this, we designed two protospacer oligos (sgRNA-7 and sgRNA-8) 196 bp apart to simultaneously target a small region in the gene Pp3c16_8300 (Figure 4A). Genotyping revealed that two plants contained apparent deletions larger than 100 bp (*n* = 31) (Figure 4A, gel). To investigate further, we sequenced these plants (plants #4 and #9) as well as three other plants exhibiting a similar band size to wild type (plants #5, #6, and #7) (Figure 4A). Interestingly, of the plants with visible deletions, plant #4 was solely targeted by sgRNA-8 and resulted in a 101 bp deletion (Figure 4A, gene models). The other deletion mutant, plant #9, was targeted by both sgRNA-7 and sgRNA-8 and resulted in two separate deletions of 152 bp and 10 bp, respectively. In this case, the cleavage of both target sites did not result in complete excision. Similarly, plant #5 was also targeted by both sgRNAs, though it only resulted in a total deletion of 8 bp. Plant #6 was only targeted by sgRNA-8, and plant #7 was not edited (Figure 4A, gene models).

**Figure 4.**
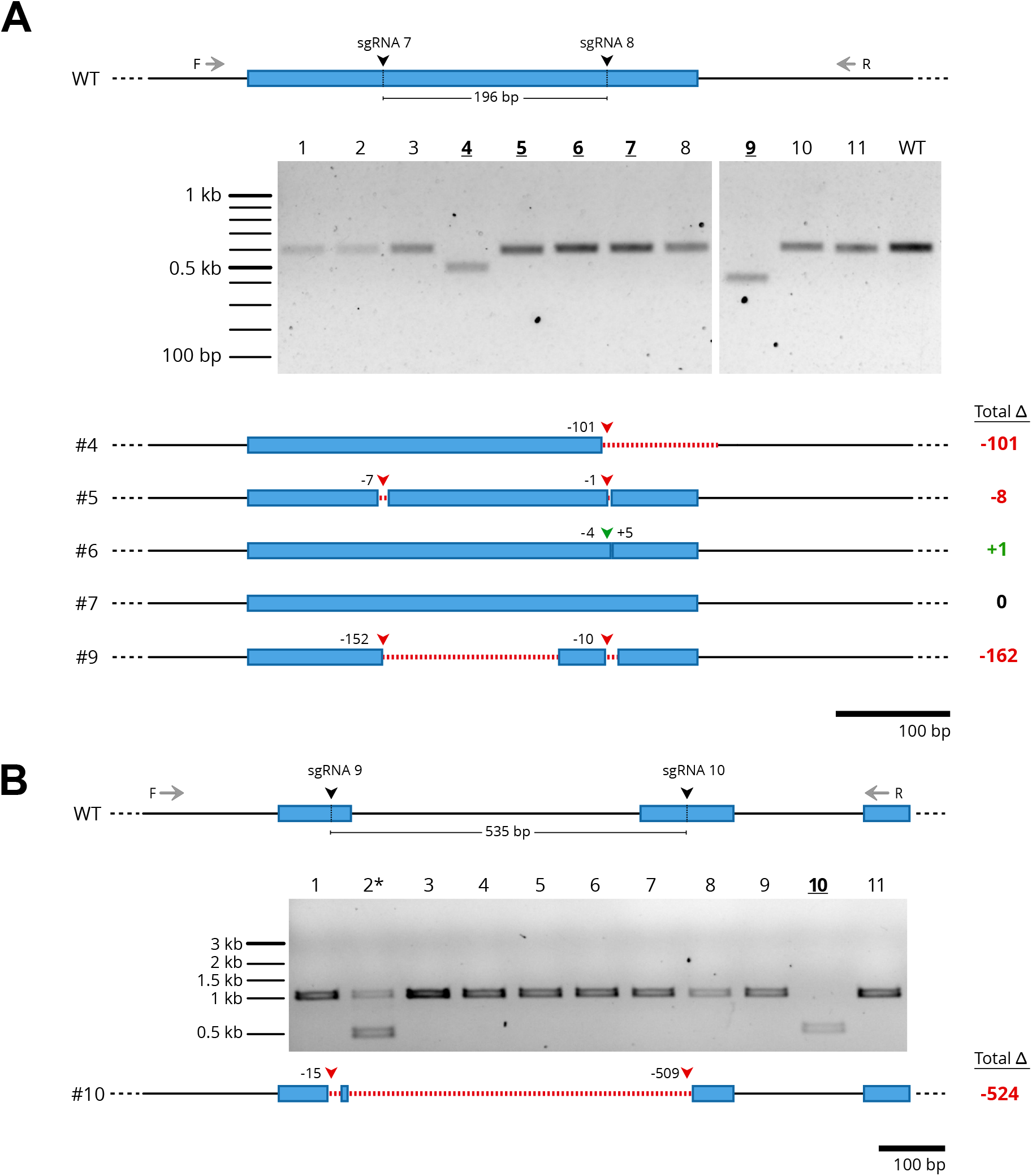
Testing genome editing efficiency in moss using two sgRNAs at adjacent sites within the same locus. The gene model of wild type (WT) and the expected sgRNA cleavage sites (196 bp apart for (A) and 535 bp apart for (B)) are displayed. Forward (F) and reverse (R) primers for genotyping are shown, with the genotyping PCR results displayed on the gel. Bolded/underlined numbers represent plants that were sequenced at the target locus. Gene models of the sequencing results are displayed below the gel with red dashed lines representing deleted regions. Red arrowheads indicate net deletion and green arrowheads represent net addition of bases at that particular sgRNA site, with the corresponding number of bases deleted or added next to each arrowhead. The total net change in base pair length for each plant is indicated to the right. The asterisk in (B) represents a chimeric plant in which the editing is suspected to have occurred after the first cell division.

We performed a similar experiment on a different gene (Pp2c9_8040) in which we designed two protospacer oligos with expected cleavage sites 535 bp apart (Figure 4B). Genotyping revealed two deletion mutants (plants #2 and #10), one of which is likely a chimeric plant due to the presence of an additional, wild type-sized fragment (plant #2) (Figure 4B, gel). Sequencing of plant #10 revealed cleavage at both sgRNA sites: a 15 bp deletion at sgRNA-9 and a 509 bp deletion at sgRNA-10. Interestingly, there remained 15 bp of wild type sequence between the two deletions, indicating that the two sgRNAs failed to excise the fragment entirely (Figure 4B, gene model). These data suggest that while it is possible to create large deletions, simultaneous targeting of adjacent sites may not increase the efficiency of uncovering knock-out mutants as both target sites may repair separately.

### Generating vectors for homology-directed repair

Homology-directed repair is an endogenous pathway that repairs DNA double-strand breaks using an available DNA template that shares regions of homology on both sides of the break site (Moynahan and Jasin 2010). Using CRISPR/Cas9, homology-directed repair can be exploited by supplying a DNA donor template together with the Cas9 enzyme and the sgRNA to repair the targeted region with extreme accuracy. To test for Cas9-induced homology-directed repair in moss, we designed a strategy to insert sequences encoding for mEGFP (Vidali et al. 2009) at the 3’ end of Pp3c22_1100 (Figure 5A). We identified a protospacer sequence that spanned the junction between the coding region and the 3’ UTR of Pp3c22_1100, which was an ideal site to target cleavage by Cas9. We generated entry clones containing homology fragments upstream and downstream of the desired insertion site (Figure 5B) and subsequently recombined them to generate the DNA donor vector containing the 5’ homology, mEGFP, and 3’ homology fragments (Figure 5C). We co-transformed the Cas9/sgRNA co-expression vector and the DNA donor vector into protoplasts. Genotyping revealed that 6 plants (28.6%) contained fragments of similar size expected for mEGFP insertion (*n* = 21, Figure 5D). Sequencing of one plant revealed seamless insertion of an in-frame mEGFP flanked by the expected attB4 and attB3 sites. Due to the high rate of successful insertions, we constructed several second-fragment entry vectors for use with three-fragment Multisite Gateway recombination for fluorescent protein gene tagging experiments (Figure 5E). These include genes encoding mEGFP (1X, 2X, and 3X) and mRuby2 (1X, 2X, and 3X) with or without DNA encoding for stop codons for C- and N-terminal protein tagging, respectively.

**Figure 5.**
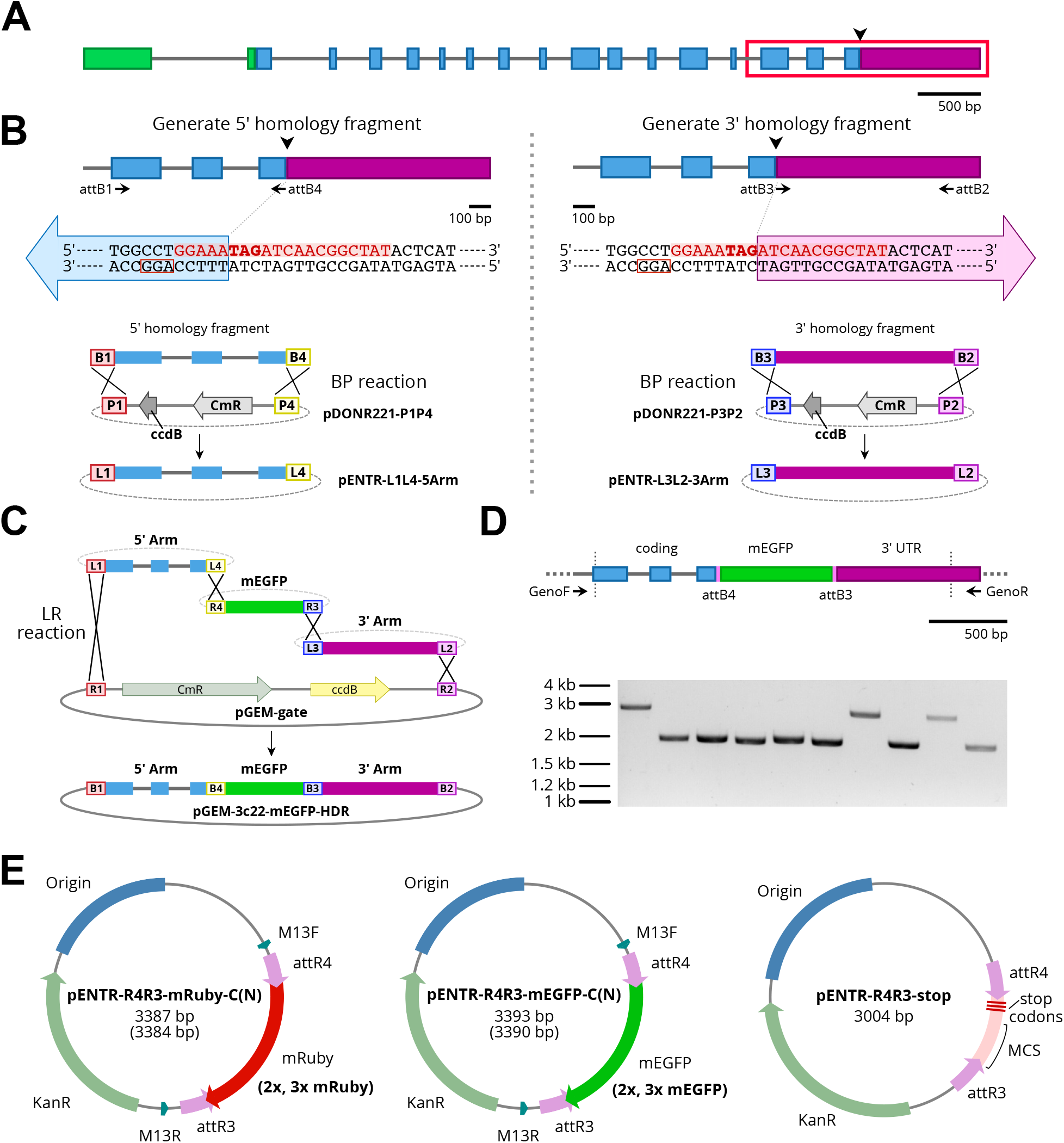
Cloning strategy for insertion of sequences encoding for mEGFP at the 3’ end of the Pp3c22_1100 gene. (A) The gene model representing the Pp3c22_1100 gene. The arrowhead represents the desired mEGFP insertion site. Exons are represented with colored boxes and introns are represented as gray lines. The 5’ UTR, the coding region, and the 3’ UTR are represented by blue, green, and purple, respectively. The red outlines the section of the gene model shown in (B). (B) Generation of the 5’ and 3’ homology fragments using PCR and primers with attB overhangs and subsequent BP reaction into first- and third-element pDONR vectors, respectively. The DNA sequence of the junction between the coding region and the 3’ UTR is shown, with bolded “TAG” representing the stop codon, red sequence representing the protospacer binding site, and the red box representing the PAM. The large blue and purple arrows represent the 5’ and 3’ homology fragments, respectively. (C) A schematic showing the entry vectors created in (B) undergoing a Gateway Multisite LR reaction (Invitrogen) with a second-element mEGFP entry vector and a destination vector to make the final DNA homology donor plasmid. (D) A gene model representing the genotyping strategy with forward and reverse primers. The region between the dashed, vertical lines represents the region present in the homology donor vector. The genotyping results are displayed on the gel. Larger bands show insertion of mEGFP into the Pp3c22_1100 locus. (E) Second-element entry vectors constructed to facilitate insertion of sequences encoding fluorescent proteins and the stop cassette using homology-directed repair.

We reasoned that homology-directed repair would also provide an ideal method to generate reliable and clean knockout alleles by, for example, inserting a fragment with an in-frame stop codon. To do this, we constructed a plasmid encoding three stop codons in each reading frame fused to a 200 bp multiple cloning site (pENTR-R4R3-stop, Figure 5E). Insertion of this “stop cassette” allows for easy identification of knockout mutants. Additionally, the presence of a multiple cloning site following the stop codons allows the use of restriction enzymes during genotyping to further validate the insertion of the stop cassette.

### Cas9 cleaves genomic DNA in the presence of competing plasmid DNA in moss cells

To perform precise genome editing using CRISPR-induced homology-directed repair, it may not always be possible to identify a protospacer sequence with high specificity near the desired editing site. Preserving the genomic sequence between the protospacer target site and the desired editing site necessitates the presence of the protospacer sequence within the DNA donor template, which is problematic. First, we reasoned that saturating amounts of DNA donor template containing the protospacer sequence might titrate away Cas9 from cleaving the genomic site. And second, the Cas9 enzyme could potentially cleave the plasmid DNA, rendering the DNA donor template inoperable during repair.

To test if cleavage of a genomic site is possible in the presence of a DNA donor template that contains the protospacer sequence, we designed the following experiment. We generated a vector (pMK-Cas9-Hyg) expressing Cas9 and a sgRNA designed to target the hygromycin resistance gene. Transformation of pMK-Cas9-Hyg alone into a moss line containing a single copy of the hygromycin resistance gene should result in genome editing that renders the plant hygromycin sensitive. However, if a donor DNA template harboring the hygromycin resistance gene is co-transformed with pMK-Cas9-Hyg, then we would expect this template to protect the genomic site from being edited, resulting in a larger percentage of hygromycin resistant plants. Surprisingly, we did not observe significant differences between the number of plants sensitive to hygromycin with (58.5%) or without (52.4%) donor template co-transformation (n = 82 plants, Figure 6). These data indicate that the presence of a plasmid containing a protospacer binding site does not inhibit Cas9 from cleaving genomic DNA in moss cells.

**Figure 6.**
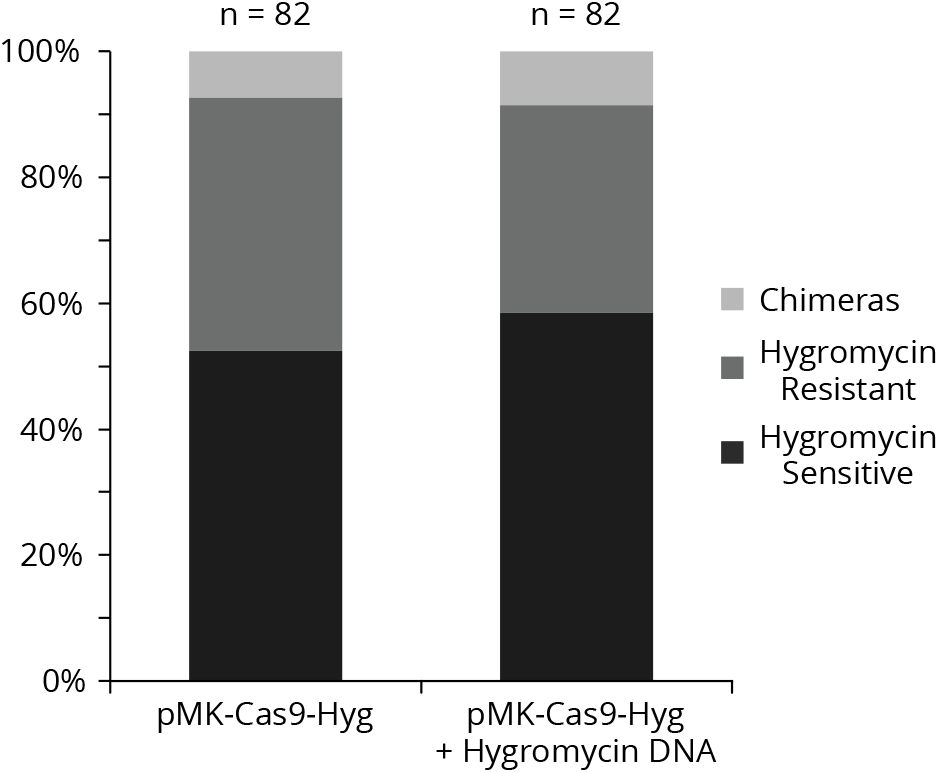
A stacked bar graph representing the amount of plants exhibiting different sensitivity to hygromycin after targeting the stable hygromycin cassette with Cas9 and with (right) or without (left) co-transformation of a plasmid containing the hygromycin sgRNA targeting sequence.

### Cas9 cleaves plasmid DNA in moss cells

Given that genomic sites were still accessible to Cas9 cleavage even in the presence of donor template DNA harboring the same target site, we wondered if Cas9 could cleave plasmid DNA in moss cells. To test this, we compared the viability of plants co-transformed with a hygromycin resistant plasmid (pGL2 as described in (Bilang et al. 1991)) and either a Cas9 plasmid targeting the hygromycin resistance gene (pMK-Cas9-Hyg) or a Cas9 plasmid targeting a non-existent site (pMK-Cas9-NGG). As expected, selection for the Cas9 plasmid resulted in similar numbers of surviving plants (733, hygromycin target; 833, nonexistent site). However, selection for the hygromycin resistant plasmid yielded strikingly different numbers (9.5, hygromycin target; 157, nonexistent site). These results show that the majority of the hygromycin resistant plasmid is cleaved by Cas9, indicating that Cas9 is able to cleave plasmid DNA that contains a protospacer sequence.

### Mutagenesis of the DNA donor template containing a protospacer sequence increases the efficiency of Cas9-induced homology directed repair

CRISPR-mediated homology-directed insertion of the mEGFP gene at the 3’ end of the Pp3c22_1100 coding region was highly successful (Figure 5). In this experiment, the protospacer sequence was at an ideal position as it extended across the end of the coding sequence to the beginning of the 3’ untranslated region and included the DNA encoding the stop codon (Figure 5B). In this instance, the protospacer sequence was not present on the DNA donor template. Rather, portions of the sequence were shared amongst the two homology fragments separated by the mEGFP coding sequence. In cases where protospacer design is suboptimal and the protospacer sequence resides on the donor DNA template, we reasoned that mutagenesis of either the protospacer sequence or the PAM in the DNA donor template should help to achieve accurate homology-directed repair. To test this, we designed a strategy to insert a gene encoding mRuby2 between the 5’ untranslated region and the beginning of the coding region of Pp3c16_8300. In this case, the protospacer targeting sequence resides within the coding region and the DNA donor plasmid, therefore, contains the protospacer sequence within the 3’ homology fragment (Figure 7A). We altered the DNA donor template to create a silent mutation within the protospacer sequence 3 bp from the PAM (5’ NGG 3’) (Figure 7B). We co-transformed protoplasts with the Cas9/sgRNA co-expression plasmid and either the mutagenized or unmutagenized DNA donor template. As expected, we were unable to detect any homology-directed repair events in plants transformed with the DNA donor template containing the protospacer sequence (n = 23 plants) (Figure 7A, gel). Conversely, homology-directed repair events were readily detected in plants transformed with mutagenized DNA donor templates: 6 out of 17 plants contained insertions with the expected size (Figure 7B). We sequenced PCR products from two of these plants and verified successful homology-directed insertions. These data demonstrate that this DNA donor template containing the protospacer sequence resulted in inefficient homology-directed repair.

**Figure 7.**
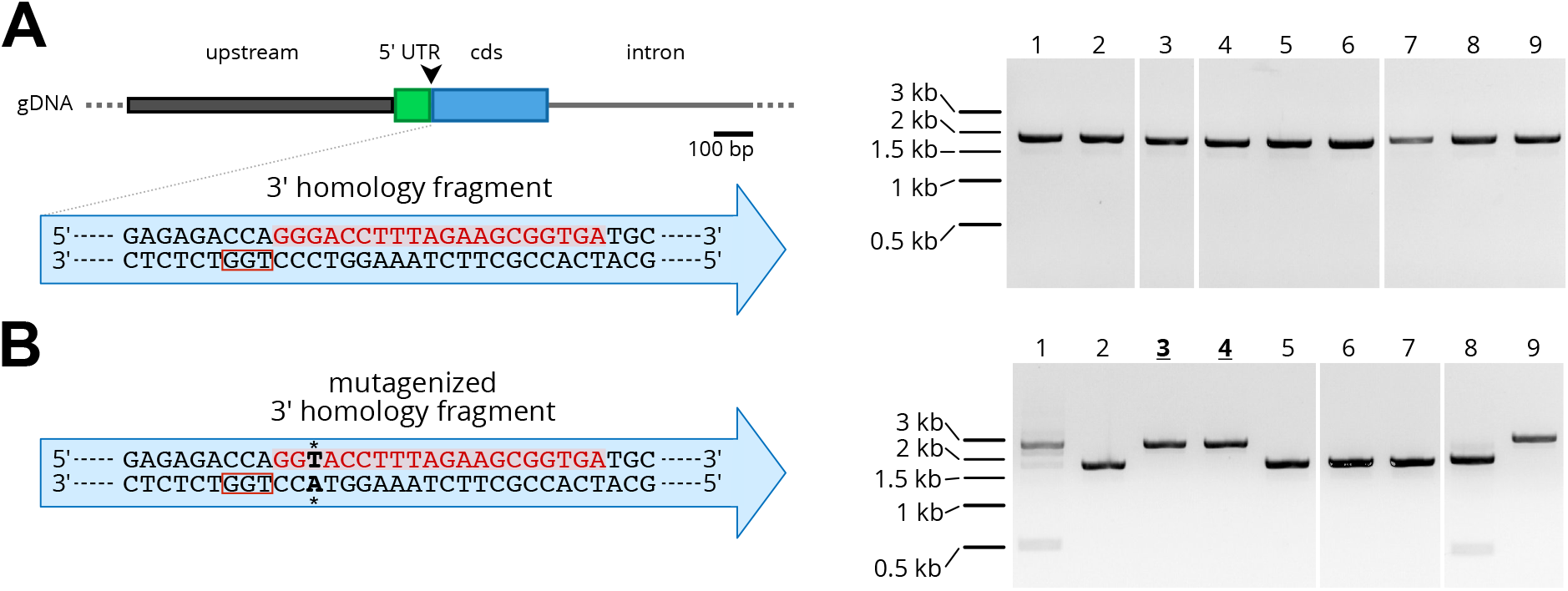
Homology-directed repair with a DNA donor homology plasmid that contains a protospacer targeting sequence. (A) A partial gene model representing the 5’ region of Pp3c16_8300. The black arrowhead represents the desired mRuby2 insertion site. The DNA sequence of the 3’ homology fragment is shown within the blue arrow, with the protospacer binding sequence shown in red and the PAM shown in a red box. Plants co-transformed with the DNA donor template and the Cas9 plasmid were unable to integrate the mRuby2 sequence into the locus (right). (B) The DNA sequence of the 3’ homology fragment after introduction of a silent mutation within the protospacer binding sequence (bold, asterisks). Mutagenesis allows mRuby2 integration upon co-transformation with the Cas9 plasmid (right). Sequenced plants are indicated with bold and underlined numbers.

## DISCUSSION

The ability to precisely edit the genome has been shown to be extremely reliable and rapid using CRISPR/Cas9 in a variety of organisms (Sander and Joung 2014; Malzahn, Lowder, and Qi 2017). Based on modifications of the vectors described by (Miao et al. 2013), we have described a CRISPR/Cas9 vector system to enhance CRISPR editing in *Physcomitrella patens* with methods that can be easily translated for use in other species. Rapid and efficient ligation of a pre-assembled, double-stranded oligo containing a custom protospacer sequence into an entry vector eliminates the need for gene synthesis. The final expression vector contains the sgRNA and the Cas9 expression cassettes. This results in equal stoichiometric amounts of DNA sequences coding for each component within each protoplast, which may be beneficial when targeting multiple genomic sites. We have also demonstrated that the native *P. patens* U6 promoter is highly efficient at expressing the sgRNA in comparison to the rice U3 promoter and therefore have included it in our vector system. Additionally, we have described an approach to rapidly generate DNA donor templates for homology-directed repair using a three-fragment Multisite Gateway reaction (Invitrogen).

As expansion of gene families is common in *Physcomitrella patens* (Zimmer et al. 2013; Lang et al. 2018), multiple family members can be targeted in a single transformation, eliminating repetitive and prolonged procedures. We have demonstrated the ability of our vector system to perform multiplex editing by targeting six different genomic sites in a single transformation using two expression vectors. Since Multisite Gateway cloning (Invitrogen) allows recombination of a maximum of four fragments, three destination vectors with different plant selection cassettes allows transformation of up to 12 sgRNAs in a single protoplast. In the test case presented here, only four of the six genomic sites were successfully edited. The absence of edits in two of the sites could be explained by differences in protospacer efficiency.

Multiplexing is not solely limited to different genes, as multiple sgRNAs can be used to target a single region. This has been shown to be successful in creating gene deletions as well as large-scale chromosomal deletions (Cai et al. 2018; Mali et al. 2013; He et al. 2015; Xiao et al. 2013; Hao et al. 2016). In our case, we tested the ability to create easy-to-detect knock-out mutations using two sgRNAs separated by ∼200 and ∼500 base pairs in protein-coding genes. Indeed, visible deletions were detected in both cases, but at relatively low frequencies. Additionally, we were unable to obtain complete removal of the intervening region which may result from differences in protospacer efficiency. It’s possible that one Cas9:sgRNA complex cleaved one site more than the other site, and/or that one site repaired before the other site was cleaved.

Induction of homology-directed repair by Cas9-mediated double-strand breaks has been shown to be highly successful in *P. patens* (Collonnier et al. 2017). Traditionally, homology-directed repair has been used in *P. patens* for decades in which a linear template containing regions of homology flanking a selectable marker is transformed into protoplasts and is integrated into the genome (Kamisugi and Cuming 2009; Prigge and Bezanilla 2010). In this case, stable integration of a selectable marker is used to isolate recombinant plants. However, integration of a selectable marker can limit genetic manipulation: N-terminal protein tagging at an endogenous gene locus is challenging due to the placement of the selection cassette, and in other instances, loxP “scar” sequences remain after successful removal of the selection cassette upon expression of the Cre recombinase (Sander and Joung 2014). In the latter case, it should be noted that removal of the selection cassette requires transient expression of Cre and therefore takes substantially longer to obtain the desired modifications. CRISPR-induced homology-directed repair is a way to combat these issues in which the selection cassette is transiently expressed from the Cas9-sgRNA expression vector. Thus, there is no need to remove it from the genome, opening the ability make extremely precise alterations to the genome.

CRISPR-induced homology-directed repair in plants has been successful using ssDNA oligos (Shan et al. 2013; Svitashev et al. 2015, 2016), linear dsDNA (Sun et al. 2016; Schiml, Fauser, and Puchta 2014), and circular (Svitashev et al. 2015; Čermák et al. 2015; J.-F. Li et al. 2013; Butler et al. 2016; Gil-Humanes et al. 2017) DNA donor templates. Here, we demonstrated the ability to perform highly efficient gene tagging of Pp3c22_1100 using a circular donor template constructed with Multisite Gateway (Invitrogen). Due to the high rate of success, we constructed a variety of second-fragment entry vectors compatible with three-fragment Multisite Gateway recombination (Invitrogen) for rapid cloning of fluorescent protein-tagging constructs (Figure 5E). These flexible, modular vectors provide a stream-lined system to rapidly generate DNA donor plasmids that will enable tagging the same gene with single, double, triple, red or green fluorescent proteins.

Homology-directed repair experiments are not solely limited to gene knock-in, however. Because moss plants can be propagated asexually, it is possible to use homology-directed repair to rapidly generate the same allele in different genetic backgrounds without the need to perform genetic crosses. Additionally, homology-directed repair can be used to create clean and easy-to-detect knock-out mutants: we constructed a “stop cassette” entry vector containing a multiple cloning site with stop codons in each reading frame. In this way, knock-out mutants can be readily detected by a simple shift in band size. Generating knock-out mutants solely with Cas9 and sgRNA is feasible, but may be less efficient due to the possibility that the surrounding microhomology could favor in-frame repair (Bae et al. 2014; Chang et al. 2017). Nevertheless, this latter approach may yield more variability in the different mutations created which could be beneficial in determining gene function. Thus, having all of these options available when performing CRISPR-mediated knock-out or knock-in experiments gives us the ability to have tight control over the desired results.

A careful choice of protospacer is essential for successful homology-directed repair. Insertion of the mEGFP coding sequence between the Pp3c22_1100 coding and 3’ UTR regions did not contain the protospacer sequence on the DNA donor template. In this scenario, the protospacer was divided in two portions on the DNA donor template separated by the mEGFP coding sequence. In cases where this particular design is not possible, we performed a series of experiments to test whether Cas9 cleaves in cases where the protospacer sequence remains on the donor template. We validated that Cas9 cleaves plasmid DNA that contains a protospacer sequence in moss cells, rendering the DNA donor template useless during repair. We also demonstrated that, while large amounts of DNA that contain the protospacer sequence are present within the cell, Cas9 retains the ability to cleave genomic DNA in moss cells. We have shown that these complications can be avoided by introducing a point mutation within the protospacer sequence present on the DNA donor template. In this case, a silent mutation 3 bases from the PAM restored successful homology-directed repair events (Figure 7B).

In conclusion, the CRISPR/Cas9 vector system presented here represents a powerful toolkit for rapid and efficient genome editing in the model moss *Physcomitrella patens*. Through non-homologous end joining, CRISPR/Cas9 permits the study of essential genes by creating potential hypomorphs as well as of nonessential genes by creating allelic variants. Through homology-directed repair, sequences can be precisely inserted or removed with co-transformation of a DNA donor template constructed *in vitro*. The simple and modular design of our vector system allows fast and economical vector construction. Additionally, editing of large gene families in a single, transient transformation is achievable in a short timeframe. Furthermore, due to the modular design, our CRISPR/Cas9 vector system could readily be employed in other organisms with slight species-specific modifications to promoter sequences.

## MATERIALS & METHODS

### Protospacer sequence design and ligation

For each editing experiment, entry vectors were linearized with *BsaI*. The CRISPOR online software (crispor.tefor.net) (Haeussler et al. 2016) was used to design protospacers for each editing experiment using *P. patens* (Phytozome V11) and *S. pyogenes* (5’ NGG 3’) as the genome and PAM parameters, respectively. Protospacers were chosen based on high specificity scores and low off-target frequency. The chosen protospacer for a given gene and its reverse complement were then constructed to have 4 nucleotides added to their 5’ ends such that, when annealed, they create sticky ends compatible with *BsaI*-linearized entry vectors (Figure 2B). These were synthesized as oligonucleotides (Supplemental Primer Table) and annealed together using a PCR machine (500 pmol of each, 10 µL total volume with PCR machine setting: 98 °C for 3 min, 0.1 °C/sec to oligo Tm, hold 10 min, 0.1 °C/sec to 25 °C). The final product was ligated into *BsaI*-linearized entry vector using Instant Sticky-End Ligation Master Mix (New England Biolabs) following the manufacturer’s recommendations.

### Polymerase III promoter assay

To build a CRISPR/Cas9 vector system for *P. patens*, we wanted to first assess RNA polymerase III promoter efficiency. The NLS-4 moss line contains a transgene that codes for nuclear-localized GFP fused to GUS (NLS-GFP-GUS) as described in (Bezanilla, Pan, and Quatrano 2003). To target NLS-GFP-GUS using a rice U3 promoter, we ligated protospacer oligos into pENTR-OsU3-sgRNA (a gift from Devin O’Connor) to create pENTR-OsU3-sgRNA-NGG. To target NLS-GFP-GUS using a *P. patens* U6 promoter, we removed the OsU3 promoter from pENTR-OsU3-sgRNA-NGG using an *AscI* and *SalI* digest. We subsequently amplified the PpU6 promoter from wild type *P. patens* Gransden strain using primers with *AscI* and *SalI* sites (Supplemental Primer Table), digested with *AscI* and *SalI*, and ligated into linearized pENTR-OsU3-sgRNA-NGG using Sticky-End Ligation Master Mix (New England Biolabs) following the manufacturer’s recommendations. These entry vectors were recombined with pH-Ubi-Cas9 (Miao et al. 2013) using an LR clonase reaction to create the final expression constructs, pH-Ubi-Cas9-OsU3-NGG and pH-Ubi-Cas9-PpU6-NGG.

Prior to imaging, we removed the labels from the plates such that the images were acquired by a blinded observer and drew a grid on the bottom of the plates to act as guides for counting. We counted 7-day old plants and recorded the presence or absence of nuclear fluorescence for each plant using a fluorescence stereomicroscope (Leica MZ16FA), equipped with the following filter: excitation 480/40, dichroic 505 long pass, emission 510 long pass.

### U6 promoter/sgRNA entry vector constructs

To generate pENTR-PpU6-sgRNA-L1L2, the first Gateway entry vector for the *P. patens* vector system, we amplified the PpU6 and sgRNA fragments with two separate PCR reactions. For the PpU6 fragment, we used a forward primer containing an *AscI* site and a reverse primer containing two inverted *BsaI* sites at the 5’ ends (Supplementary Primer Table). Similarly, for the sgRNA fragment we used a forward primer containing two inverted *BsaI* sites and a reverse primer containing a *SalI* site (Supplementary Primer Table). The two fragments were then ligated using an overlap extension PCR reaction and ligated into pGEM/T-Easy (Promega). Positive clones were digested with *AscI* and *SalI* and the dropout was subsequently ligated into an *AscI*- and *SalI*-linearized pENTR-PpU6-sgRNA-NGG plasmid.

To generate the six entry vectors compatible with Multisite Gateway (Invitrogen) for multiplexing experiments, we amplified the sgRNA expression cassette from pENTR-PpU6-sgRNA-L1L2 using primers (Supplemental Primer Table) containing different Multisite Gateway attachment sites (attB) and subsequently recombined with the Multisite Gateway pDONR221 plasmid set (Invitrogen) using a BP clonase reaction following the manufacturer’s recommendations.

### Cas9/sgRNA destination and expression constructs

To generate the three destination vectors, we purified a fragment containing the Cas9 and Gateway cassette from pH-Ubi-Cas9 (Miao et al. 2013) digested with *StuI* and *PmeI*. This fragment was ligated into linearized pMH, pMK, and pZeo vectors by blunt-end ligation to create pMH-Cas9-gate, pMK-Cas9-gate, and pZeo-Cas9-gate. Sequences are available on AddGene (https://www.addgene.org/kits/bezanilla-crispr-physcomitrella/). All of the Cas9/sgRNA expression vectors used in this study were generated using Gateway (for one sgRNA) or Multisite Gateway (for multiple sgRNAs) to recombine the entry vectors and destination vectors just described (Invitrogen).

### Homology-directed repair constructs

To generate pENTR-R4R3-stop (the “stop cassette”), we amplified 360 bp of the plasmid pBluescriptSK(+), including the multiple cloning site, with attB4r and attB3r Gateway primers (Supplemental Primer Table). The forward primer also contained three stop codons in each frame. We subsequently cloned the PCR fragment into pDONR221-P4rP3r (Invitrogen) using a BP clonase reaction following the manufacturer’s recommendations.

To generate the mEGFP and mRuby2 tagging entry vectors, we amplified mEGFP (Vidali et al. 2009) and mRuby2 (Lam et al. 2012) coding sequences using forward and reverse primers (Supplemental Primer Table) that contained attB4r and attB3r sites, respectively. The forward primers (Supplemental Primer Table) for both mEGFP and mRuby2 also contained a *BamHI* site. These PCR products were subsequently cloned into pDONR221-P4rP3r (Invitrogen) using a BP clonase reaction to create pENTR-R4R3-mEGFP-C and pENTR-R4R3-mRuby-C. mEGFP-pGEM, a vector described by (Vidali et al. 2009), contains *BamHI* and *BglII* sites flanking the mEGFP coding sequence. We generated mRuby2-pGEM, a vector constructed in the same way as mEGFP-pGEM (Vidali et al. 2009). We digested these vectors with *BamHI* and *BglII* and the resulting fragments were ligated into *BamHI*-digested pENTR-R4R3-mEGFP-C and pENTR-R4R3-mRuby-C to create pENTR-R4R3-2XmEGFP-C and pENTR-R4R3-2XmRuby-C, respectively. We linearized the resulting 2X constructs with *BamHI* and ligated the *BamHI*/*BglII* fragments from mEGFP-pGEM and mRuby2-pGEM to create the 3X constructs, pENTR-R4R3-3XmEGFP-C and pENTR-R4R3-3XmRuby-C, respectively. To create the mEGFP and mRuby2 N-terminal constructs, the process above was repeated except the attB3r primers (Supplemental Primer Table) did not contain stop codons.

### DNA donor templates

We used the three-fragment Multisite Gateway cloning system (Invitrogen) to generate the final homology-directed repair homology constructs. For Pp3c22_1100, we amplified 2 fragments of approximately 800 bp upstream and downstream of the Pp3c22_1100 stop codon. For Pp3c16_8300, we amplified 2 fragments of approximately 800 bp upstream and downstream of the expected start codon. For both genes, we cloned the upstream fragments into pDONR221-P1P4 and the downstream fragments into pDONR221-P3P2 using a BP clonase reaction. To create the final homology-directed repair DNA donor plasmids, the resulting pENTR vectors from the BP reaction underwent an LR clonase reaction with the second-position tagging vector (pENTR-R4R3-mEGFP-C for Pp3c22_1100 and pENTR-R4R3-mRuby-N for Pp3c16_8300) and the destination vector, pGEM-gate (Vidali et al. 2009).

To restore efficient homology-directed repair while tagging Pp3c16_8300, we performed site-directed mutagenesis on the entry vector containing the 3’ homology fragment (pENTR-L3L2-3c16-3Arm) to create pENTR-L3L2-3c16-3Arm-mut by altering the third nucleotide from the PAM (5’ NGG 3’) within the protospacer. We repeated the LR reaction using this third-position entry vector to generate pGEM-3c16-mRuby-HDR-mut.

### Moss tissue culture and transformation

We propagated moss tissue weekly by light homogenization and subsequently plated on 10-cm petri dishes to maintain the protonemal growth stage. Dishes contained 25 mL PpNH4 growth medium (103 mM MgSO_4_, 1.86 mM KH_2_PO_4_, 3.3 mM Ca(NO_3_)_2_, 2.72 mM (NH_4_)_2_-tartrate, 45 µM FeSO_4_, 9.93 µM H_3_BO_3_, 220 nM CuSO_4_, 1.966 µM MnCl_2_, 231 nM CoCl_2_, 191 nM ZnSO_4_, 169 nM KI, and 103 nM Na_2_MoO_4_) with 0.7% agar covered with cellophane disks. Plants were grown in daily cycles of 16 h light/8 h dark with 85 µmol photons m^−2^ s^−1^. For transformation, protoplasts were transformed with 30 µg of each DNA construct using PEG-mediated transformation protocol (as described in (Augustine et al. 2011)). Plants were allowed to regenerate on plant regeneration media (PRMB) for 4 days (described in (Wu and Bezanilla 2014)) atop of cellophane disks. Depending upon the selection cassette present on the expression vector, we subsequently moved plants to PpNH4 growth media containing either hygromycin (15 µg/mL), G418 (20 µg/mL), or Zeocin (50 µg/mL). Plants were not selected for homology-directed repair DNA donor vectors. After 7 days on selection, we moved plants to PpNH4 media without antibiotics for maximal growth. Plants were allowed to grow for 2-3 weeks until tufts were 0.5-1 cm in diameter for DNA extraction.

### DNA extraction and genotyping

We extracted DNA from plants that were 3-4 weeks old (0.5-1cm in diameter) using the protocol as described in (Augustine et al. 2011). For editing experiments, we used PCR primers (Supplemental Primer Table) surrounding the expected Cas9 cleavage site (∼300-400 bp on each side). For homology-directed repair experiments, we used PCR primers outside of the homology region to avoid amplification of residual DNA donor template. To perform PCR, we used Q5 polymerase (New England Biolabs) using the manufacturer’s recommendations.

### T7 endonuclease assay

To detect CRISPR edits, we amplified a 0.5 to 1 kb genomic region flanking the potential CRISPR editing site by PCR. The PCR product from each candidate plant was mixed with a roughly equivalent amount of wild type PCR product of the same locus. The mixture was denatured and annealed in a PCR machine and subsequently digested with 1 µL of T7 endonuclease (New England Biolabs) following the manufacturer’s recommendations. We examined the digest on a 1% agarose gel.

## Supporting information

Supplemental Primer Table

## SUPPLEMENTAL DATA

**Table S1 –** A table of primers and oligos used in this study.

**Figure S1**

**Figure S2**

## ACKNOWLEDGEMENTS

We thank Fabien Nogué for the *P. patens* U6 promoter sequence. M.C. received support from the Plant Biology Graduate Program at the University of Massachusetts, Amherst and X.C. received support from the Molecular and Cellular Biology Graduate Program at Dartmouth College. Additionally, this work was supported by NSF grants MCB-1330171 and MCB-1715785 to M.B.

## AUTHOR CONTRIBUTIONS

D.R.M., M.C., X.C., and M.B. designed the research. D.R.M., M.C., and X.C. performed the research. D.R.M. and M.B. wrote the article.

## LITERATURE CITED

Augustine, R. C., K. A. Pattavina, E. Tuzel, L. Vidali, and M. Bezanilla. 2011. “Actin Interacting Protein1 and Actin Depolymerizing Factor Drive Rapid Actin Dynamics in Physcomitrella Patens.” The Plant Cell 23 (10): 3696–3710. https://doi.org/10.1105/tpc.111.090753.

Bae, Sangsu, Jiyeon Kweon, Heon Seok Kim, and Jin-Soo Kim. 2014. “Microhomology-Based Choice of Cas9 Nuclease Target Sites.” Nature Methods 11 (7): 705–6. https://doi.org/10.1038/nmeth.3015.

Beucher, Andrea, Julie Birraux, Leopoldine Tchouandong, Olivia Barton, Atsushi Shibata, Sandro Conrad, Aaron A Goodarzi, Andrea Krempler, Penny A Jeggo, and Markus Löbrich. 2009. “ATM and Artemis Promote Homologous Recombination of Radiation-Induced DNA Double-Strand Breaks in G2.” The EMBO Journal 28 (21): 3413–27. https://doi.org/10.1038/emboj.2009.276.

Bezanilla, M., Aihong Pan, and Ralph S. Quatrano. 2003. “RNA Interference in the Moss Physcomitrella Patens.” PLANT PHYSIOLOGY 133 (2): 470–74. https://doi.org/10.1104/pp.103.024901.

Bilang, Roland, Shigeru Iida, Alex Peterhans, Ingo Potrykus, and Jerzy Paszkowski. 1991. “The 3′-Terminal Region of the Hygromycin-B-Resistance Gene Is Important for Its Activity in Escherichia Coli and Nicotiana Tabacum.” Gene 100 (April): 247–50. https://doi.org/10.1016/0378-1119(91)90375-L.

Butler, Nathaniel M., Nicholas J. Baltes, Daniel F. Voytas, and David S. Douches. 2016. “Geminivirus-Mediated Genome Editing in Potato (Solanum Tuberosum L.) Using Sequence-Specific Nucleases.” Frontiers in Plant Science 7 (July). https://doi.org/10.3389/fpls.2016.01045.

Cai, Yupeng, Li Chen, Shi Sun, Cunxiang Wu, Weiwei Yao, Bingjun Jiang, Tianfu Han, and Wensheng Hou. 2018. “CRISPR/Cas9-Mediated Deletion of Large Genomic Fragments in Soybean.” International Journal of Molecular Sciences 19 (12): 3835. https://doi.org/10.3390/ijms19123835.

Čermák, Tomáš, Nicholas J. Baltes, Radim Čegan, Yong Zhang, and Daniel F. Voytas. 2015. “High-Frequency, Precise Modification of the Tomato Genome.” Genome Biology 16 (1): 232. https://doi.org/10.1186/s13059-015-0796-9.

Chang, Howard H. Y., Nicholas R. Pannunzio, Noritaka Adachi, and Michael R. Lieber. 2017. “Non-Homologous DNA End Joining and Alternative Pathways to Double-Strand Break Repair.” Nature Reviews Molecular Cell Biology 18 (8): 495–506. https://doi.org/10.1038/nrm.2017.48.

Chen, Xiugui, Xuke Lu, Na Shu, Shuai Wang, Junjuan Wang, Delong Wang, Lixue Guo, and Wuwei Ye. 2017. “Targeted Mutagenesis in Cotton (Gossypium Hirsutum L.) Using the CRISPR/Cas9 System.” Scientific Reports 7 (1): 44304. https://doi.org/10.1038/srep44304.

Collonnier, Cécile, Aline Epert, Kostlend Mara, François Maclot, Anouchka Guyon-Debast, Florence Charlot, Charles White, Didier G. Schaefer, and Fabien Nogué. 2017. “CRISPR-Cas9-Mediated Efficient Directed Mutagenesis and RAD51-Dependent and RAD51-Independent Gene Targeting in the Moss *Physcomitrella Patens*.” Plant Biotechnology Journal 15 (1): 122–31. https://doi.org/10.1111/pbi.12596.

Feng, Z., Y. Mao, N. Xu, B. Zhang, P. Wei, D.-L. Yang, Z. Wang, et al. 2014. “Multigeneration Analysis Reveals the Inheritance, Specificity, and Patterns of CRISPR/Cas-Induced Gene Modifications in Arabidopsis.” Proceedings of the National Academy of Sciences 111 (12): 4632–37. https://doi.org/10.1073/pnas.1400822111.

Gil-Humanes, Javier, Yanpeng Wang, Zhen Liang, Qiwei Shan, Carmen V. Ozuna, Susana Sánchez-León, Nicholas J. Baltes, et al. 2017. “High-Efficiency Gene Targeting in Hexaploid Wheat Using DNA Replicons and CRISPR/Cas9.” The Plant Journal 89 (6): 1251–62. https://doi.org/10.1111/tpj.13446.

Haeussler, Maximilian, Kai Schönig, Hélène Eckert, Alexis Eschstruth, Joffrey Mianné, Jean-Baptiste Renaud, Sylvie Schneider-Maunoury, et al. 2016. “Evaluation of Off-Target and on-Target Scoring Algorithms and Integration into the Guide RNA Selection Tool CRISPOR.” Genome Biology 17 (1): 148. https://doi.org/10.1186/s13059-016-1012-2.

Hao, Huanhuan, Xiaofei Wang, Haiyan Jia, Miao Yu, Xiaoyu Zhang, Hui Tang, and Liping Zhang. 2016. “Large Fragment Deletion Using a CRISPR/Cas9 System in Saccharomyces Cerevisiae.” Analytical Biochemistry 509 (September): 118–23. https://doi.org/10.1016/j.ab.2016.07.008.

He, Zuyong, Chris Proudfoot, Alan J. Mileham, David G. McLaren, C. Bruce A. Whitelaw, and Simon G. Lillico. 2015. “Highly Efficient Targeted Chromosome Deletions Using CRISPR/Cas9: Highly Efficient Targeted Chromosome Deletions.” Biotechnology and Bioengineering 112 (5): 1060–64. https://doi.org/10.1002/bit.25490.

Jinek, M., K. Chylinski, I. Fonfara, M. Hauer, J. A. Doudna, and E. Charpentier. 2012. “A Programmable Dual-RNA-Guided DNA Endonuclease in Adaptive Bacterial Immunity.” Science 337 (6096): 816–21. https://doi.org/10.1126/science.1225829.

Kamisugi, Yasuko, and Andrew C. Cuming. 2009. “Gene Targeting.” In The Moss Physcomitrella Patens, edited by Celia D. Knight, Pierre-Franois Perroud, and David J. Cove, 76–112. Oxford, UK: Wiley-Blackwell. https://doi.org/10.1002/9781444316070.ch4.

Lam, Amy J, François St-Pierre, Yiyang Gong, Jesse D Marshall, Paula J Cranfill, Michelle A Baird, Michael R McKeown, et al. 2012. “Improving FRET Dynamic Range with Bright Green and Red Fluorescent Proteins.” Nature Methods 9 (10): 1005–12. https://doi.org/10.1038/nmeth.2171.

Lang, Daniel, Kristian K. Ullrich, Florent Murat, Jörg Fuchs, Jerry Jenkins, Fabian B. Haas, Mathieu Piednoel, et al. 2018. “The *Physcomitrella Patens* Chromosome-Scale Assembly Reveals Moss Genome Structure and Evolution.” The Plant Journal 93 (3): 515–33. https://doi.org/10.1111/tpj.13801.

Li, Chao, Turgay Unver, and Baohong Zhang. 2017. “A High-Efficiency CRISPR/Cas9 System for Targeted Mutagenesis in Cotton (Gossypium Hirsutum L.).” Scientific Reports 7 (1): 43902. https://doi.org/10.1038/srep43902.

Li, Jian-Feng, Julie E Norville, John Aach, Matthew McCormack, Dandan Zhang, Jenifer Bush, George M Church, and Jen Sheen. 2013. “Multiplex and Homologous Recombination–Mediated Genome Editing in Arabidopsis and Nicotiana Benthamiana Using Guide RNA and Cas9.” Nature Biotechnology 31 (8): 688–91. https://doi.org/10.1038/nbt.2654.

Li, Meiru, Xiaoxia Li, Zejiao Zhou, Pingzhi Wu, Maichun Fang, Xiaoping Pan, Qiupeng Lin, Wanbin Luo, Guojiang Wu, and Hongqing Li. 2016. “Reassessment of the Four Yield-Related Genes Gn1a, DEP1, GS3, and IPA1 in Rice Using a CRISPR/Cas9 System.” Frontiers in Plant Science 7 (March). https://doi.org/10.3389/fpls.2016.00377.

Liang, Zhen, Kang Zhang, Kunling Chen, and Caixia Gao. 2014. “Targeted Mutagenesis in Zea Mays Using TALENs and the CRISPR/Cas System.” Journal of Genetics and Genomics 41 (2): 63–68. https://doi.org/10.1016/j.jgg.2013.12.001.

Lopez-Obando, Mauricio, Beate Hoffmann, Carine Géry, Anouchka Guyon-Debast, Evelyne Téoulé, Catherine Rameau, Sandrine Bonhomme, and Fabien Nogué. 2016. “Simple and Efficient Targeting of Multiple Genes Through CRISPR-Cas9 in *Physcomitrella Patens*.” G3: Genes|Genomes|Genetics 6 (11): 3647–53. https://doi.org/10.1534/g3.116.033266.

Makarova, Kira S., Daniel H. Haft, Rodolphe Barrangou, Stan J. J. Brouns, Emmanuelle Charpentier, Philippe Horvath, Sylvain Moineau, et al. 2011. “Evolution and Classification of the CRISPR–Cas Systems.” Nature Reviews Microbiology 9 (6): 467–77. https://doi.org/10.1038/nrmicro2577.

Mali, P., L. Yang, K. M. Esvelt, J. Aach, M. Guell, J. E. DiCarlo, J. E. Norville, and G. M. Church. 2013. “RNA-Guided Human Genome Engineering via Cas9.” Science 339 (6121): 823–26. https://doi.org/10.1126/science.1232033.

Malzahn, Aimee, Levi Lowder, and Yiping Qi. 2017. “Plant Genome Editing with TALEN and CRISPR.” Cell & Bioscience 7 (1): 21. https://doi.org/10.1186/s13578-017-0148-4.

Miao, Jin, Dongshu Guo, Jinzhe Zhang, Qingpei Huang, Genji Qin, Xin Zhang, Jianmin Wan, Hongya Gu, and Li-Jia Qu. 2013. “Targeted Mutagenesis in Rice Using CRISPR-Cas System.” Cell Research 23 (10): 1233–36. https://doi.org/10.1038/cr.2013.123.

Moynahan, Mary Ellen, and Maria Jasin. 2010. “Mitotic Homologous Recombination Maintains Genomic Stability and Suppresses Tumorigenesis.” Nature Reviews Molecular Cell Biology 11 (3): 196–207. https://doi.org/10.1038/nrm2851.

Nishimasu, Hiroshi, F. Ann Ran, Patrick D. Hsu, Silvana Konermann, Soraya I. Shehata, Naoshi Dohmae, Ryuichiro Ishitani, Feng Zhang, and Osamu Nureki. 2014. “Crystal Structure of Cas9 in Complex with Guide RNA and Target DNA.” Cell 156 (5): 935–49. https://doi.org/10.1016/j.cell.2014.02.001.

Prigge, M. J., and M. Bezanilla. 2010. “Evolutionary Crossroads in Developmental Biology: Physcomitrella Patens.” Development 137 (21): 3535–43. https://doi.org/10.1242/dev.049023.

Puchta, H. 2005. “The Repair of Double-Strand Breaks in Plants: Mechanisms and Consequences for Genome Evolution.” Journal of Experimental Botany. https://doi.org/10.1093/jxb/eri025.

Pyott, Douglas E., Emma Sheehan, and Attila Molnar. 2016. “Engineering of CRISPR/Cas9-Mediated Potyvirus Resistance in Transgene-Free *Arabidopsis* Plants: CRISPR/Cas9-Induced Potyvirus Resistance.” Molecular Plant Pathology 17 (8): 1276–88. https://doi.org/10.1111/mpp.12417.

Saidi, Younousse, Andrija Finka, Mickhail Chakhporanian, Jean-Pierre Zrÿd, Didier G. Schaefer, and Pierre Goloubinoff. 2005. “Controlled Expression of Recombinant Proteins in Physcomitrella Patens by a Conditional Heat-Shock Promoter: A Tool for Plant Research and Biotechnology.” Plant Molecular Biology 59 (5): 697–711. https://doi.org/10.1007/s11103-005-0889-z.

Sander, Jeffry D, and J Keith Joung. 2014. “CRISPR-Cas Systems for Editing, Regulating and Targeting Genomes.” Nature Biotechnology 32 (4): 347–55. https://doi.org/10.1038/nbt.2842.

Sargent, R G, M A Brenneman, and J H Wilson. 1997. “Repair of Site-Specific Double-Strand Breaks in a Mammalian Chromosome by Homologous and Illegitimate Recombination.” Molecular and Cellular Biology 17 (1): 267–77. https://doi.org/10.1128/MCB.17.1.267.

Schaefer, Didier G., and Jean-Pierre Zryd. 1997. “Efficient Gene Targeting in the Moss Physcomitrella Patens.” The Plant Journal 11 (6): 1195–1206. https://doi.org/10.1046/j.1365-313X.1997.11061195.x.

Schiml, Simon, Friedrich Fauser, and Holger Puchta. 2014. “The CRISPR/Cas System Can Be Used as Nuclease for *in Planta* Gene Targeting and as Paired Nickases for Directed Mutagenesis in Arabidopsis Resulting in Heritable Progeny.” The Plant Journal 80 (6): 1139–50. https://doi.org/10.1111/tpj.12704.

Shan, Qiwei, Yanpeng Wang, Jun Li, Yi Zhang, Kunling Chen, Zhen Liang, Kang Zhang, et al. 2013. “Targeted Genome Modification of Crop Plants Using a CRISPR-Cas System.” Nature Biotechnology 31 (8): 686–88. https://doi.org/10.1038/nbt.2650.

Shi, Jinrui, Huirong Gao, Hongyu Wang, H. Renee Lafitte, Rayeann L. Archibald, Meizhu Yang, Salim M. Hakimi, Hua Mo, and Jeffrey E. Habben. 2017. “ARGOS8 Variants Generated by CRISPR-Cas9 Improve Maize Grain Yield under Field Drought Stress Conditions.” Plant Biotechnology Journal 15 (2): 207–16. https://doi.org/10.1111/pbi.12603.

Sun, Yongwei, Xin Zhang, Chuanyin Wu, Yubing He, Youzhi Ma, Han Hou, Xiuping Guo, Wenming Du, Yunde Zhao, and Lanqin Xia. 2016. “Engineering Herbicide-Resistant Rice Plants through CRISPR/Cas9-Mediated Homologous Recombination of Acetolactate Synthase.” Molecular Plant 9 (4): 628–31. https://doi.org/10.1016/j.molp.2016.01.001.

Svitashev, Sergei, Christine Schwartz, Brian Lenderts, Joshua K. Young, and A. Mark Cigan. 2016. “Genome Editing in Maize Directed by CRISPR–Cas9 Ribonucleoprotein Complexes.” Nature Communications 7 (1): 13274. https://doi.org/10.1038/ncomms13274.

Svitashev, Sergei, Joshua K. Young, Christine Schwartz, Huirong Gao, S. Carl Falco, and A. Mark Cigan. 2015. “Targeted Mutagenesis, Precise Gene Editing, and Site-Specific Gene Insertion in Maize Using Cas9 and Guide RNA.” Plant Physiology 169 (2): 931–45. https://doi.org/10.1104/pp.15.00793.

Vidali, L., P. A. C. van Gisbergen, C. Guerin, P. Franco, M. Li, G. M. Burkart, R. C. Augustine, L. Blanchoin, and M. Bezanilla. 2009. “Rapid Formin-Mediated Actin-Filament Elongation Is Essential for Polarized Plant Cell Growth.” Proceedings of the National Academy of Sciences 106 (32): 13341–46. https://doi.org/10.1073/pnas.0901170106.

Wang, Fujun, Chunlian Wang, Piqing Liu, Cailin Lei, Wei Hao, Ying Gao, Yao-Guang Liu, and Kaijun Zhao. 2016. “Enhanced Rice Blast Resistance by CRISPR/Cas9-Targeted Mutagenesis of the ERF Transcription Factor Gene OsERF922.” Edited by Richard A Wilson. PLOS ONE 11 (4): e0154027. https://doi.org/10.1371/journal.pone.0154027.

Wilson, Fiona M., Kate Harrison, Andrew D. Armitage, Andrew J. Simkin, and Richard J. Harrison. 2019. “CRISPR/Cas9-Mediated Mutagenesis of Phytoene Desaturase in Diploid and Octoploid Strawberry.” Plant Methods 15 (1): 45. https://doi.org/10.1186/s13007-019-0428-6.

Wu, Shu-Zon, and Magdalena Bezanilla. 2014. “Myosin VIII Associates with Microtubule Ends and Together with Actin Plays a Role in Guiding Plant Cell Division.” ELife 3 (September): e03498. https://doi.org/10.7554/eLife.03498.

Xiao, An, Zhanxiang Wang, Yingying Hu, Yingdan Wu, Zhou Luo, Zhipeng Yang, Yao Zu, et al. 2013. “Chromosomal Deletions and Inversions Mediated by TALENs and CRISPR/Cas in Zebrafish.” Nucleic Acids Research 41 (14): e141–e141. https://doi.org/10.1093/nar/gkt464.

Zhang, Yi, Zhen Liang, Yuan Zong, Yanpeng Wang, Jinxing Liu, Kunling Chen, Jin-Long Qiu, and Caixia Gao. 2016. “Efficient and Transgene-Free Genome Editing in Wheat through Transient Expression of CRISPR/Cas9 DNA or RNA.” Nature Communications 7 (1): 12617. https://doi.org/10.1038/ncomms12617.

Zimmer, Andreas D, Daniel Lang, Karol Buchta, Stephane Rombauts, Tomoaki Nishiyama, Mitsuyasu Hasebe, Yves Van de Peer, Stefan A Rensing, and Ralf Reski. 2013. “Reannotation and Extended Community Resources for the Genome of the Non-Seed Plant Physcomitrella Patens Provide Insights into the Evolution of Plant Gene Structures and Functions.” BMC Genomics 14 (1): 498. https://doi.org/10.1186/1471-2164-14-498.

