## Supplemental Primer Table for "Efficient and Modular CRISPR-Cas9 Vector System for *Physcomitrella patens*"

**Table S1** – A table of primers and oligos used in this study

| Primer ID | Sequence (5'->3') | Purpose |
|---|---|---|
| UM_2149 | GGCGCGCCGAGCTCGAATTCGTCCATTGAAGC | PpU6 promoter amplicon |
| UM_2151 | GTCGACCCTGCCATGGGTTGTAAGTCCTCCAC CTTCC | PpU6 promoter amplicon |
| UM_2206 | TGAGACCTTGGTCTCTATGGGTTGTAAGTCCT CCACCTTC | PpU6 promoter + BsaI amplicon |
| UM_2207 | AGAGACCAAGGTCTCAGTTTTAGAGCTATGC | BsaI + sgRNA amplicon |
| UM_2208 | CAGTCACGACGTTGTAAAACGACGG | BsaI + sgRNA amplicon |
| UM_2320 | GGGGACAAGTTTGTACAAAAAAGCAGGCTCCC CTTCACCGTCAGATGC | pENTR-PpU6P-L1L2-sgRNA, pENTR-PpU6P-L1R5-sgRNA, pENTR-PpU6P-L1L4-sgRNA insert |
| UM_2321 | GGGGACAACTTTTGTATACAAAGTTGTGAGCT CGAATTCGTCCATTGAAGC | pENTR-PpU6P-L1R5-sgRNA insert |
| UM_2322 | GGGGACAACTTTGTATACAAAAGTTGCCCCTT CACCGTCAGATGC | pENTR-PpU6P-L5L2-sgRNA, pENTR-PpU6P-L5L4-sgRNA insert |
| UM_2323 | GGGGACCACTTTGTACAAGAAAGCTGGGTAGA GCTCGAATTCGTCCATTGAAGC | pENTR-PpU6P-L1L2-sgRNA, pENTR-PpU6P-L3L2-sgRNA, pENTR-PpU6P-L5L2-sgRNA insert |
| UM_2324 | GGGGACAACTTTGTATAGAAAAGTTGGGTGGA GCTCGAATTCGTCCATTGAAGC | pENTR-PpU6P-L1L4-sgRNA, pENTR-PpU6P-L5L4-sgRNA insert |
| UM_2325 | GGGGACAACTTTTCTATACAAAGTTGCCCCTT CACCGTCAGATGC | pENTR-PpU6P-R4R3-sgRNA insert |
| UM_2326 | GGGGACAACTTTATTATACAAAGTTGTGAGCT CGAATTCGTCCATTGAAGC | pENTR-PpU6P-R4R3-sgRNA insert |
| UM_2327 | GGGGACAACTTTGTATAATAAAGTTGCCCCTT CACCGTCAGATGC | pENTR-PpU6P-L3L2-sgRNA insert |
| DC_173 | GGGGACAACTTTTCTATACAAAGTTGTAGCTG ATTAAGTTGGGTAACGCCAGG | stop cassette amplicon |
| DC_174 | GGGGACAACTTTATTATACAAAGTTGTAGGCT TTACACTTTATGCTTCCG | stop cassette amplicon |
| UM_2117 | GGCAGGGTCGACAATGGTGAGCAA | NLS-GFP-GUS protospacer |
| UM_2118 | AAACTTGCTCACCATTGTCGACCC | NLS-GFP-GUS protospacer |
| UM_2657 | CCATCTATCTTCGAACGCATGCGG | sgRNA-1 protospacer (Pp3c8_18830V3.1) |
| UM_2658 | AAACCCGCATGCGTTCGAAGATAG | sgRNA-1 protospacer (Pp3c8_18830V3.1) |
| UM_2695 | TCCATCCGTCCTCAGAAG | Site 1 genotyping amplicon (Pp3c8_18830V3.1) |
| UM_2696 | AACGACTATCAAATGGTGCC | Site 1 genotyping amplicon (Pp3c8_18830V3.1) |
| UM_2659 | CCATCACAGGTGCAGTCCCGTGTG | sgRNA-5 protospacer (Pp3c8_18850V3) |
| UM_2660 | AAACCACACGGGACTGCACCTGTG | sgRNA-5 protospacer (Pp3c8_18850V3) |
| UM_2697 | CTACTTAGGGCACAGCTTC | Site 5 genotyping amplicon (Pp3c8_18850V3) |
| UM_2698 | CTACTAGGTGGTCAGCAAC | Site 5 genotyping amplicon (Pp3c8_18850V3) |
| UM_2661 | CCATCTGCGGGAGAACAGACAGAA | sgRNA-6 protospacer (Pp3c23_15670V3) |
| UM_2662 | AAACTTCTGTCTGTTCTCCCGCAG | sgRNA-6 protospacer (Pp3c23_15670V3) |
| UM_2699 | TGCTGTAGGTTGCTTGAG | Site 6 genotyping amplicon (Pp3c23_15670V3) |
| UM_2700 | CTGAAATTGCCTTACTTCGC | Site 6 genotyping amplicon (Pp3c23_15670V3) |
| UM_2663 | CCATTTATGCCGACGGTCCAGTGA | sgRNA-2 protospacer (Pp3c18_4770V3) |

| | | |
|---|---|---|
| UM_2664 | AAACTCACTGGACCGTCGGCATAA | sgRNA-2 protospacer (Pp3c18_4770V3) |
| UM_2701 | AGCTAAAGCTGTTGGACG | Site 2 genotyping amplicon (Pp3c18_4770V3) |
| UM_2702 | AGCCTTCTTGTACGCATTC | Site 2 genotyping amplicon (Pp3c18_4770V3) |
| UM_2665 | CCATGCTATGACCAACGTCCCCTC | sgRNA-3 protospacer (Pp3c22_15110V3) |
| UM_2666 | AAACGAGGGGACGTTGGTCATAGC | sgRNA-3 protospacer (Pp3c22_15110V3) |
| UM_2703 | TATTCTACGATATGCAGTTGGG | Site 3 genotyping amplicon (Pp3c22_15110V3) |
| UM_2704 | CCTGAGTCCTTCTTCCTG | Site 3 genotyping amplicon (Pp3c22_15110V3) |
| UM_2655 | CCATGACAGCCCTCGTAATGCCGA | sgRNA-4 protospacer (Pp3c4_16430V3) |
| UM_2656 | AAACTCGGCATTACGAGGGCTGTC | sgRNA-4 protospacer (Pp3c4_16430V3) |
| UM_2693 | ATTGACAAGGTGGGATGTG | Site 4 genotyping amplicon (Pp3c4_16430V3) |
| UM_2694 | GAGTCCAGACAAACTTCAGC | Site 4 genotyping amplicon (Pp3c4_16430V3) |
| DC_9 | CCATTCACCGCTTCTAAAGGTCCC | sgRNA-7 protospacer (Pp3c16_8300) |
| DC_10 | AAACGGGACCTTTAGAAGCGGTGA | sgRNA-7 protospacer (Pp3c16_8300) |
| DC_11 | CCATCACGCGAACCAGATTAATCA | sgRNA-8 protospacer (Pp3c16_8300) |
| DC_12 | AAACTGATTAATCTGGTTCGCGTG | sgRNA-8 protospacer (Pp3c16_8300) |
| DC_45 | CTTCGTCATGGCGCAGG | Pp3c16_8300 genotyping amplicon |
| DC_461 | GCCCAGCTTCAAACGTAACC | Pp3c16_8300 genotyping amplicon |
| DC_30 | CCATAGCACAGGCGACGGTTGCAC | sgRNA-9 protospacer (Pp2c9_8040) |
| DC_31 | AAACGTGCAACCGTCGCCTGTGCT | sgRNA-9 protospacer (Pp2c9_8040) |
| DC_32 | CCATCGGCTTTGCTTGGAATAGAC | sgRNA-10 protospacer (Pp2c9_8040) |
| DC_33 | AAACGTCTATTCCAAGCAAAGCCG | sgRNA-10 protospacer (Pp2c9_8040) |
| DC_61 | TCACCGTCCTCGTTGTAG | Pp2c9_8040 genotyping amplicon |
| DC_62 | CAAACATCGACAAGCTACACG | Pp2c9_8040 genotyping amplicon |
| DC_118 | CCATATAGCCGTTGATCTATTTCC | Pp3c22_1100 protospacer |
| DC_119 | AAACGGAAATAGATCAACGGCTAT | Pp3c22_1100 protospacer |
| DC_108 | GGGGACAAGTTTGTACAAAAAAGCAGGCTACT ACAAGCGGAGATTGCTCAAGC | Pp3c22_1100 5' homology amplicon |
| DC_109 | GGGGACAACTTTGTATAGAAAAGTTGGGTGTT TCCAGGCCACAGCATCAATCTC | Pp3c22_1100 5' homology amplicon |
| DC_112 | GGGGACAACTTTGTATAATAAAGTTGATCAAC GGCTATACTCATTTCTCC | Pp3c22_1100 3' homology amplicon |
| DC_113 | GGGGACCACTTTGTACAAGAAAGCTGGGTAAT CTCACCTTCCGTTCTAAATACG | Pp3c22_1100 3' homology amplicon |
| DC_120 | CATTTCGTCGGAGTGTCGG | Pp3c22_1100 genotyping amplicon |
| DC_121 | GGTGTCACAGGTTGATGAAGC | Pp3c22_1100 genotyping amplicon |
| DC_225 | CCATCGATTCCTTGCGGTCCGAAT | Hygromycin resistance protospacer |
| DC_226 | AAACATTCGGACCGCAAGGAATCG | Hygromycin resistance protospacer |
| DC_9 | CCATTCACCGCTTCTAAAGGTCCC | Pp3c16_8300 tagging protospacer |
| DC_10 | AAACGGGACCTTTAGAAGCGGTGA | Pp3c16_8300 tagging protospacer |
| DC_179 | GGGGACAAGTTTGTACAAAAAAGCAGGCTCAT GGAAACATGGTTGAAATTATACG | Pp3c16_8300 5' homology amplicon |
| DC_180 | GGGGACAACTTTGTATAGAAAAGTTGGGTGAG AAACGCAAATCGACAGC | Pp3c16_8300 5' homology amplicon |

| | | |
|---|---|---|
| DC_181 | GGGGACAACTTTGTATAATAAAGTTGGGATGTATGTCAGAGAGAGACCAGG | Pp3c16_8300 3' homology amplicon |
| DC_182 | GGGGACCACTTTGTACAAGAAAGCTGGGTGAAAGTATAATCACAGGCCTCAACG | Pp3c16_8300 3' homology amplicon |
| DC_461 | CCTTTCCACAATTTAAATTTAAAGAATAAGACATGAG | Pp3c16_8300 HDR genotying amplicon |
| DC_401 | GCACGAACATGAGTGAGG | Pp3c16_8300 HDR genotying amplicon |
| DC_272 | GACCAGGTACCTTTAGAAGC | Pp3c16_8300 site-directed mutagenesis |
| DC_273 | GCTTCTAAAGGTACCTGGTC | Pp3c16_8300 site-directed mutagenesis |
| DC_159 | GGGGACAACTTTTCTATACAAAGTTGAAGGATCCATGGTGAGCAAGGGCGAG | mEGFP and mRuby2 cds amplicon (C+N term) |
| DC_111 | GGGGACAACTTTATTATACAAAGTTGTTTACTTGTACAGCTCGTCCATGC | mEGFP and mRuby2 cds amplicon (C term) |
| DC_178 | GGGGACAACTTTATTATACAAAGTTGTCTTGTACAGCTCGTCCATGCC | mEGFP and mRuby2 cds amplicon (N-term) |
| EYFPF | AATGGGATCCATGGTGAGCAAGGGCGAG | mRuby-pGEM insert |
| EYFPR | AATGAGATCTCTTGTACAGCTCGTCCATGC | mRuby-pGEM insert |
