## Supplemental Figure 1 for "Efficient and Modular CRISPR-Cas9 Vector System for *Physcomitrella patens*"

```
NLS fragment                            GFP fragment
TCTTGTGCCAGTTTCGGCCGGGTCGACAATGGTGAGCAAGGGCGAGGAGCTGTTCACCGG  Wild type
TCTTGTGCCAGTTTCGGCCGGGTCGACAATGGTGAG----------GAGCTGTTCACCGG  10 bp deletion (n = 5)
TCTTGTGCCAGTTTCGGCCGGGTCGACAA----------GGGCGAGGAGCTGTTCACCGG  10 bp deletion (n = 4)
TCTTGTGCCAGTTTCGGCCGGGTCGACAATGGTGAGGAAGGGCGAGGAGCTGTTCACCGG  1 bp substitution (n = 1)
```

**Supplemental Figure 1**
Sequencing data of a subset of NLS-4 plants (*n* = 10) expressing a sgRNA targeting NLS-GFP-GUS with the OsU3 promoter. The red region represents the sgRNA binding site and the black arrow represents the expected Cas9 cleavage site.
