## Supplemental Figure 2 for "Efficient and Modular CRISPR-Cas9 Vector System for *Physcomitrella patens*"

### Site 3

| Line # | Sequence | |
|---|---|---|
| WT | GCCCTCTGCATCATCTCCGGAGGGGACGTTGGTCATAGCCAAAACTCAATC | |
| 2 | GCCCTCTGCATCATCTCCG---------TTGGTCATAGCCAAAACTCAATC | 9 bp deletion |
| 3, 22 | GCCCTCTGCATCATCTCCGGA-----CGTTGGTCATAGCCAAAACTCAATC | 5 bp deletion |
| 13 | GCCCTCTGCATCATCTCCGG---GGACGTTGGTCATAGCCAAAACTCAATC | 3 bp deletion |
| 23 | GCCCTCTGCATCATCTCCGGA----ACGTTGGTCATAGCCAAAACTCAATC | 4 bp deletion |

### Site 4

| Line # | Sequence | |
|---|---|---|
| WT | CGGTAATGGATTACCCTCG----GCATTACGAGGGCTGTCCTGTTTTATC |  |
| 2, 23 | CGGTAATGGATTACCCTCGTAATGCATTACGAGGGCTGTCCTGTTTTATC | 4 bp insertion |
| 3 | CGGTAATGGATTACCCTCG----GTATGAC-AGGGCTGTCCTGTTTTATC | 1 bp deletion, 2 bp substitution |
| 13 | CGGTAATA---------------GCATTACGAGGGCTGTCCTGTTTTATC | 11 bp deletion, 1 bp substitution |
| 20 | CGGTAATGGATTACCC--------CATTACGAGGGCTGTCCTGTTTTATC | 4 bp deletion |
| 22 | CGGTAATGGATTACCCTCG-----CATTACGAGGGCTGTCCTGTTTTATC | 1 bp deletion |

### Site 5

| Line # | Sequence | |
|---|---|---|
| WT | CTCAACAGGAGAGTCCACACACGGGACTGCACCTGTGAAGGCAGATGATG | |
| 2 | CTCAACAGGAGAGTC--CACACGGGACTGCACCTGTGAAGGCAGATGATG | 2 bp deletion |
| 3, 22 | CTCAACAGGA----------------CTGCACCTGTGAAGGCAGATGATG | 16 bp deletion |
| 13, 23 | CTCAACAGGAGAGTCCACAC------------CTGTGAAGGCAGATGATG | 12 bp deletion |
| 20 | CTCAACAGGAGAGTCCACAC--GGGACTGCACCTGTGAAGGCAGATGATG | 2 bp deletion |

### Site 6

| Line # | Sequence | |
|---|---|---|
| WT | CTTCACTCTCTCTGCGGGAGAACAGACAGAATGGATTTCGTGCTTT | |
| 2 | CTTCACTCTCTCTGCGGGAGAACAGACAGAATGGATTTCGTGCTTT | no edits |
| 3 | CTTCACTCTCTCTGCGGGAGAACAAACGGAATGGATTTCGTGCTTT | 2 bp substitution |
| 13, 20, 22, 23 | CTTCACTCTCTCTGCGGGAGAACAGA----ATGGATTTCGTGCTTT | 4 bp deletion |

**Supplemental Figure 2**
Sequence data displaying the results of the multiplex targeting experiment. The highlighted red text represents the protospacer and the black arrow represents the expected Cas9 cleavage site. Dashed lines indicate deletions and/or insertions. A summary of the genotyping data is shown in Figure 3B.
